# The BioGenome Portal: a web-based platform for biodiversity genomics data management

**DOI:** 10.1101/2023.12.20.572408

**Authors:** Emilio Righi, Roderic Guigó

## Abstract

Biodiversity genomics projects are underway with the aim of sequencing the genomes of all eukaryotic species on Earth. Here we describe the BioGenome Portal, a web-based application to facilitate organization and access to the data produced by biodiversity genomics projects. The portal integrates user-generated data with data deposited in public repositories. The portal generates sequence status reports that can be eventually ingested by designated meta-data tracking systems, facilitating the coordination task of these systems. The portal is open-source and fully customizable. It can be deployed at any site with minimum effort, contributing to the democratization of biodiversity genomics projects. Here, we illustrate the features of the BioGenome Portal through two specific instances. One instance corresponds to the Earth Biogenome Project, the worldwide umbrella for most biodiversity genomics projects. The other instance corresponds to the Catalan Initiative for the Earth Biogenome Project, a regional project aiming to sequencing the genomes of the species of the Catalan Linguistic Area.

## INTRODUCTION

Biodiversity genomics projects are underway which aim to sequence the genomes of the approximately two million known eukaryotic species on Earth. The impact of these projects will be unprecedented as it will radically alter our understanding of life on Earth. These projects differ on size and scope: some target geographic locations, such as the Darwin Tree of Life (DToL, (1) ), the Catalan Initiative for the Earth Biogenome Project (CBP, Corominas et al., submitted as a companion to NARGAB), the European Reference Genome Atlas (ERGA, (2) ) or the California Conservation Genomics Project (CCGP, (3) ), others target specific taxa, such as the vertebrate genome project (VGP, (4) ) or the 10,000 Bird Genomes (B10K, (5) ). They also differ on funding schemas, organization and structure. The Earth Biogenome Project (EBP, (6)) acts as a network umbrella to all these projects providing common guidelines and standards (operational, but also ethical and legal) for sample collection and processing, sequencing and assembling, annotation, data analysis, IT and informatics. As of 2023, the EBP includes 54 affiliated projects world-wide. Many of these projects are networks themselves, composed of multiple nodes (for instance, each European country is a node of ERGA, which includes, in addition, some regional nodes). The EBP can thus be properly described as a network of networks.

These projects (or even the nodes within a project) are operationally independent and carry out all of the steps of the genome sequencing process autonomously: from the collection of the biological samples to the production of the genome assembly. The EBP uses Genomes on a Tree (GoaT, (7)) for coordination. GoaT is a centralized resource sponsored by Tree of Life programme (https://www.sanger.ac.uk/programme/tree-of-life/) that collates observed and estimated genome-relevant metadata—including genome sizes and karyotypes—for eukaryotic species, and that also holds declarations of current and planned activity across the EBP nodes. GoaT is also the official sequence status tracker of the EBP.

Here, we describe, the BioGenome Portal (BGP), a platform that tracks, integrates and manages the data generated under a given biodiversity genomics project (not necessarily an EBP node), that helps in the coordination among the groups within the same project and, by generating a GoaT compliant sequencing status report, contributes to keep the sequencing status of the EBP up to date.

An instance of the BGP is currently used to track all the data generated under the CBP umbrella (https://dades.biogenoma.cat/) We have also deployed an instance of the BGP through which data generated by the EBP and its affiliated projects can be accessed (https://ebp.biogenoma.cat/). The BGP is highly customizable and can be also used to manage and disseminate data, which is specific to a particular site, such as vernacular names of species, and extended information about the species not available through GoaT. This could include photos, o literature text or texts emphasizing the relevance of the species within a given culture, demographic history, scientific publications, etc.).

## MATERIAL AND METHODS

### System Architecture

The BGP is composed of four Docker (8) containers orchestrated by Docker Compose. With the use of a docker-compose configuration file, it is possible to configure the application infrastructure and start all containers with a single command. This greatly simplifies application infrastructure management, improving application portability between development and production environments and facilitating continuous application integration and deployment.

### Back-end Container

The back-end container consists in a python Flask application (https://flask.palletsprojects.com ) that exposes an API via a uWSGI web server, that in turn interfaces with the NGINX (9) reverse proxy of the Front-end container, to perform CRUD (Create, Read, Update and Delete) operations against the database container.

### Front-end Container

The front-end container consists of a Vue.js ( https://vuejs.org/ ) Single Page Application (SPA) served via a NGINX reverse proxy. This container is accessible via a web browser and communicates with the back-end container via API. The SPA is composed of a Content Management System (CMS) section where authenticated users are able to manage the data present in the BGP database, and an open access section that contains many cutting-edge features to enhance data visualization.

### Cronjob Container

The cronjob container is used to make scheduled authenticated requests to the API, thus triggering specific jobs to import and update data in the database.

### Database Container

The database container is composed of a MongoDB (https://www.mongodb.com ) Docker image where all the data is stored. MongoDB uses a flexible document data model: data in the database can be stored without having to define a strict schema beforehand. In MongoDB, data is stored as documents, which are similar to JSON objects and can be easily modified and updated. One of the advantages of this flexible schema is that it allows for easy modification and expansion of the data model. As new requirements arise, users can easily add new fields to their documents without the need for complex schema migrations or downtime.

### Configuration

The BGP can be adapted to the specific needs of any biodiversity research project thanks to its configuration flexibility. Via configuration files, the BGP can be customized to import and process data from external sources and to adapt the BGP’ s charts to specific needs (**Figure 1**).

**Figure 1.**
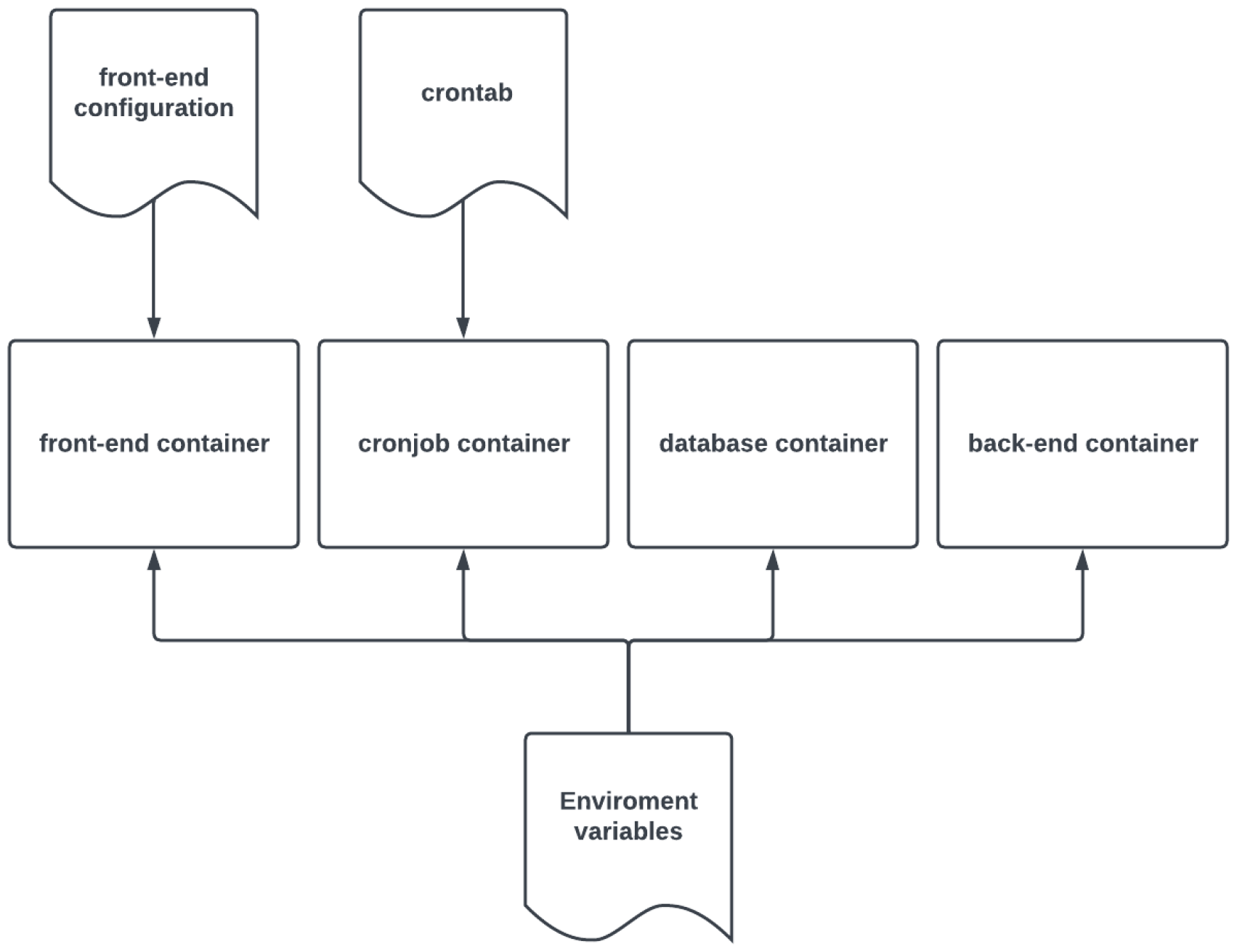
Schema depicting the configuration files injected in the BGP’ s containers

### .env

This file contains all the environment variables that will be injected into each docker container. Here it is possible to declare: the admin credentials needed to access the CMS, the taxonomic identifier to be used as the root for the phylogenetic trees present in the User-Interface (UI), the INSDC Bioproject Accession to retrieve the data from INSDC, a comma separated list of project names, to fetch sample metadata from the EBI Biosamples API, which may have not yet been associated to an assembly in the Bioproject, and other configuration variables needed to interface the docker containers between each other.

### config.json

This file allows users to personalize the look and functionality of the SPA, including the selection of charts to be displayed on each page, the ability to change the application title and logo, and the flexibility to set language preferences. By editing specific JSON entries, users can create a unique and user-centric research environment that meets their individual needs, enhances the overall user experience and provides multi-language support.

### crontab

This file is used to configure the scheduled jobs present in the crontab container.

### Data Import and Export

The BGP offers robust data management features, providing researchers a seamless way to work with biodiversity genomic data. The BGP is API-centered thus, importing and exporting data can also be done by directly querying the API.

### INSDC Data Import

The INSDC related data can be imported into the BGP database either via cronjob or CMS (**Figure 2**). Via cronjob all the data generated under the BioProject accession number declared in the .*env* file, are retrieved and stored into the database, thus enabling data tracking of that BioProject. Via CMS, authenticated users can individually import Experiments, BioSamples and Assemblies through their respective accession number.

**Figure 2.**
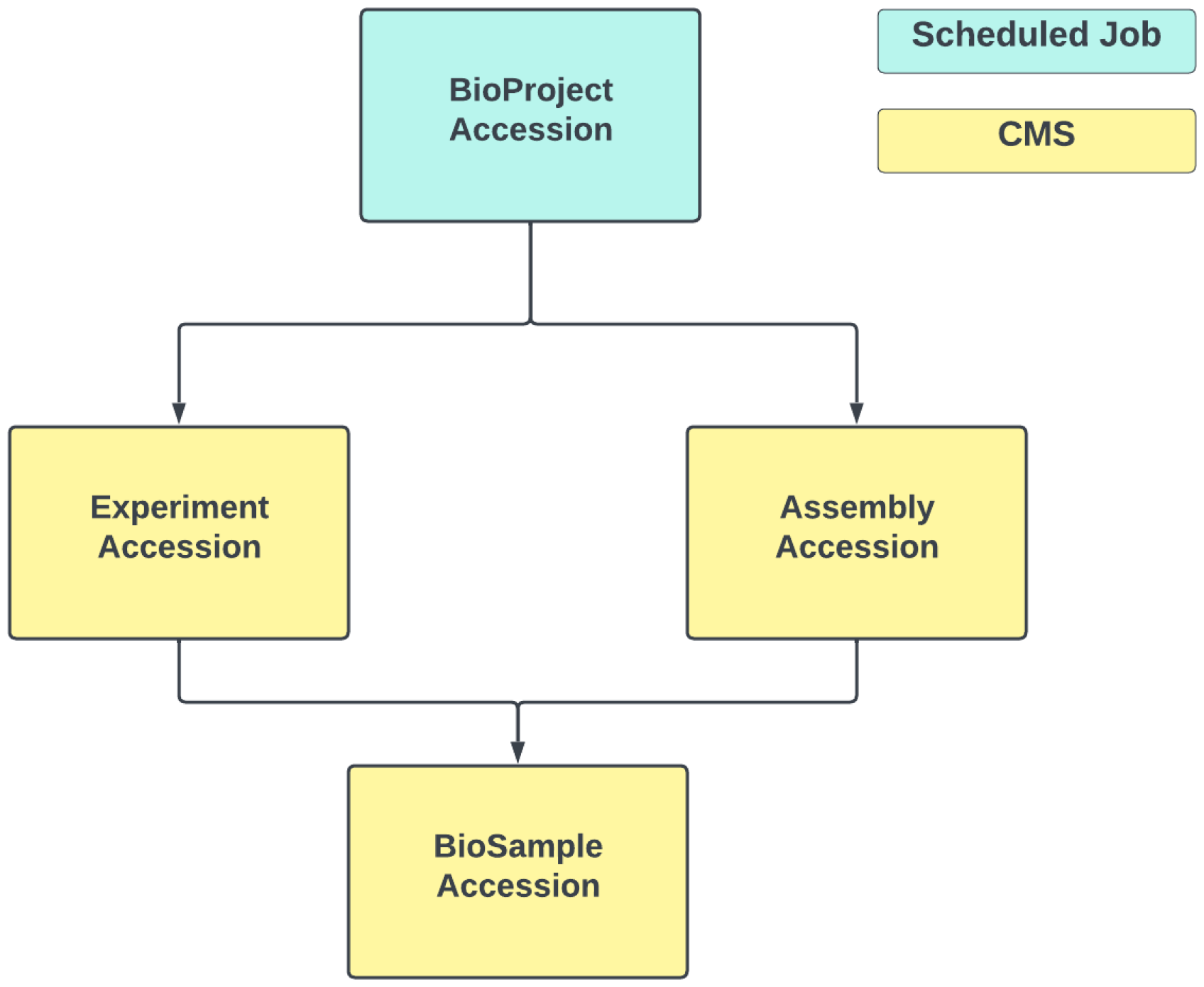
Schema depicting the INSDC data import strategies of the BGP. Via scheduled job all the INSDC data under a specific bioproject accession can be imported. Via CMS individual INSDC data can be imported via their accession.

### Samples Metadata Import

Spreadsheets containing sample metadata can be uploaded into the database via CMS. The only spreadsheet’ s columns required are the unique identifier of the sample, the scientific name and the taxonomic identifier, the remaining columns will be stored as metadata.

### GoaT Sequencing Status

The BGP offers a powerful feature that allows users to generate detailed TSV (Tab-Separated Values (TSV) files containing the sequencing status of all organisms stored in the database. This functionality is designed to provide researchers with a convenient means of aggregating and sharing important sequencing status information. The TSV file format conforms to the established GoaT sequencing report template. This alignment with the GoaT template ensures that the exported data is standardized, making it easily compatible with existing genomic data management systems and facilitating seamless integration with broader genomics research initiatives. Researchers can rely on these TSV files to generate comprehensive sequencing status reports for their data, ensuring data consistency and facilitating collaboration within the biodiversity genomics community. The sequencing status of each species can be update either automatically or manually (pre-INSDC submission)

### UI Components

The BGP has a variety of useful visualization tools such as charts, phylogenetic trees, maps (**Figure S1**) and an integrated genome browser (**Figure S2**). The charts in the BGP provide a visual representation of the data contained in the database, such as the INSDC submission status of the project, the submission trend of INSDC related data, the sequencing platform used, the genome assembly level and the habitat of the collected samples.

The phylogenetic trees allow users to explore the relationships between different organisms based on their taxonomy. By selecting a specific taxonomic parent, users can visualize the related organisms in two different types of phylogenetic trees: the indented tree and the radial tree.

The built-in maps can be used to display the organism’ s geographical distribution and the organisms collected by each country.The BGP features two types of interactive maps: a 2D map (Leaflet.js, https://leafletjs.com/) and a 3D map (CesiumJS: https://cesium.com/). With this feature, BGP makes it easy to browse and analyze the distribution of organisms in different geographic regions.

The integrated genome browser (JBrowse2 (10) ) provides users with the ability to explore DNA sequences and complete genomes using a RefGet API-compliant plugin (https://github.com/guigolab/jbrowse-plugin-refget-api). This plugin retrieves the nucleotide sequences from servers that are compliant with RefGet API (11) standards, without the need to download the entire genome assembly. In addition, users can associate genomic annotations with genomes and display those annotations alongside the corresponding genomes.

## Database Model

The BGP database model is organism-centered (**Figure 3**). At the core of this model is the association of both locally curated data and data from the INSDC with a unified set of taxonomic details. There are several important advantages to this design approach. It promotes an intuitive and structured organization of data, allowing researchers to easily navigate and retrieve information about specific organisms. It also ensures the consistency and coherence of the data representation, eliminating redundancy and potential conflicts in taxonomic classifications. In addition, an organism-centered model simplifies the process of integrating external INSDC data with locally generated information to create a consistent and integrated repository. Ultimately, this design facilitates efficient data retrieval, comparative analysis, and cross-referencing between different datasets.

**Figure 3.**
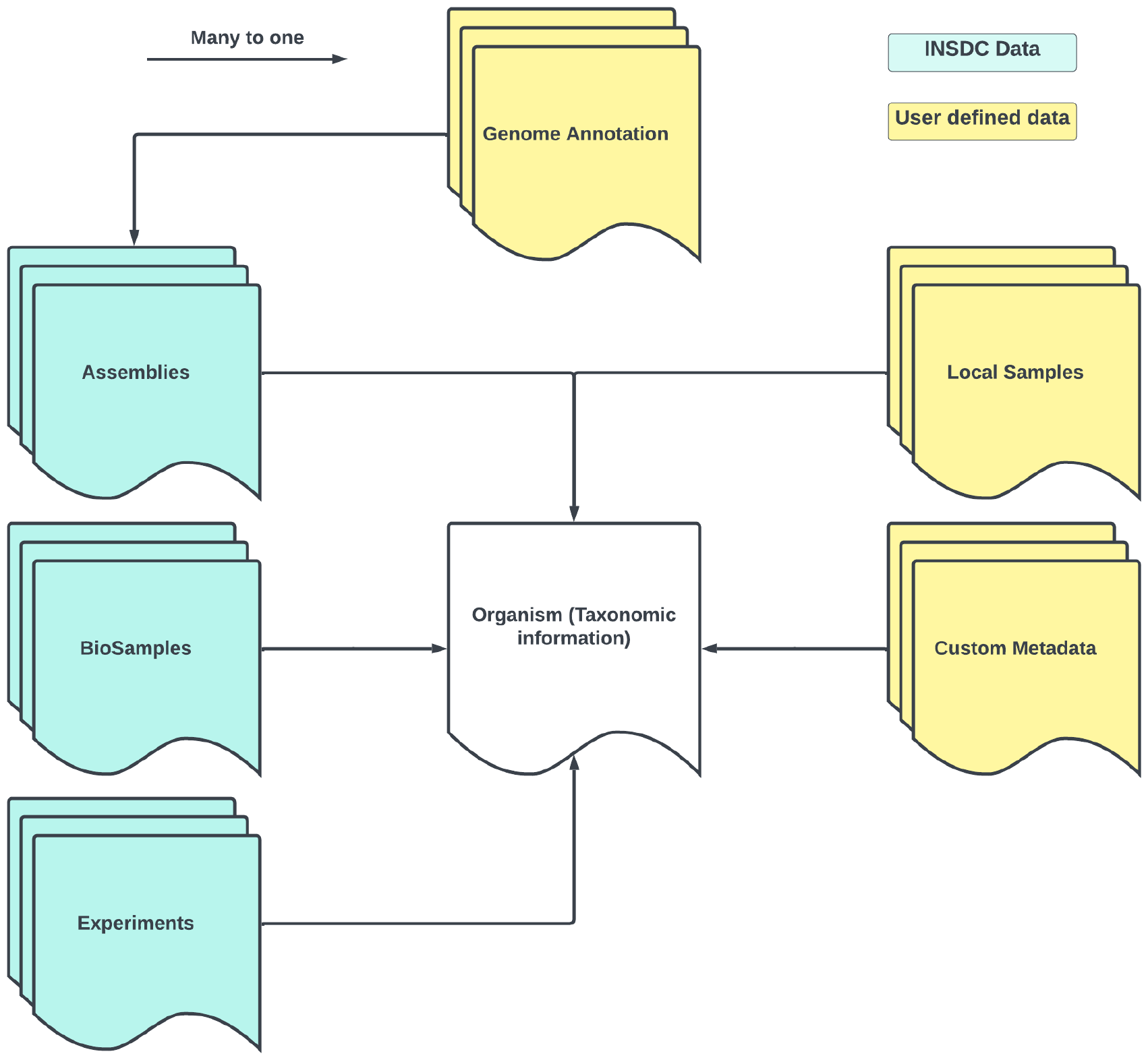
Schema depicting the organism-centered database model. In yellow all of the data that can be inserted by users. In blue the data that can be imported from INSDC via their accession number

A detailed description on how to configure, run and deploy a BGP instance can be found in the README file of the github repository (https://github.com/guigolab/biogenome-portal, DOI: 10.5281/zenodo.8314305.)

## RESULTS

We have developed the BioGenome Portal (BGP https://github.com/guigolab/biogenome-portal) to facilitate access to data produced by biodiversity genomics projects and interface public data repositories. The BGP is a web-based application that transparently brings together into a single virtual platform, all data generated under a given biodiversity project, including both data already published on INSDC, as well as data declared prior to INSDC submission. In addition, through the BGP it is possible to attach to the sequenced organisms information not available at the INSDC, such as photos, links to relevant scientific publications and other resources, historical notes, descriptions of sequenced species, species names in local languages, genome structural and functional annotations, and any other user defined information data.

The technical details of the BGP architecture are described in Material and Methods. The BGP repository can be found at https://github.com/guigolab/biogenome-portal, which also contains all the details to configure, run and deploy it Here we describe two use cases that demonstrate the adaptability, flexibility and generality of the BGP. The first use case is the BGP instance dedicated to the Catalan Initiative for the Earth BioGenome Project (CBP, Corominas et al., submitted as a companion to NARGAB). This case highlights the potentiality of the portal to supporting regional biodiversity genomics efforts. The second case is the BGP instance corresponding to the entire EBP instance. This instance showcases the capacity of the portal to deal with large amounts of data. Both together demonstrate the capacity of the BGP to provide information at different levels of resolution.

### CBP Instance

A dedicated BGP’ s instance designed to support the research objectives of the Catalan Initiative for the Earth BioGenome Project (CBP) can be found at: https://dades.biogenoma.cat/, The CBP has as a primary aim to sequence the genomes of the eukaryotic species living in the Catalan Linguistic Area (CLA). The instance tracks all the data published to INSDC under the CBP umbrella (PRJEB49670), and contains, in addition, manually curated data, such as photos, vernacular names and related publications of the sequenced organisms. This instance also provides the GoaT sequencing status report of the CBP to GoaT, thus enabling real-time updates of the CBP progress to the EBP, prior to INSDC submission.

As of December 2023, the CBP instance contains a total of 58 organisms (**Figure 4**), five of them with already published assemblies (12–16).

**Figure 4.**
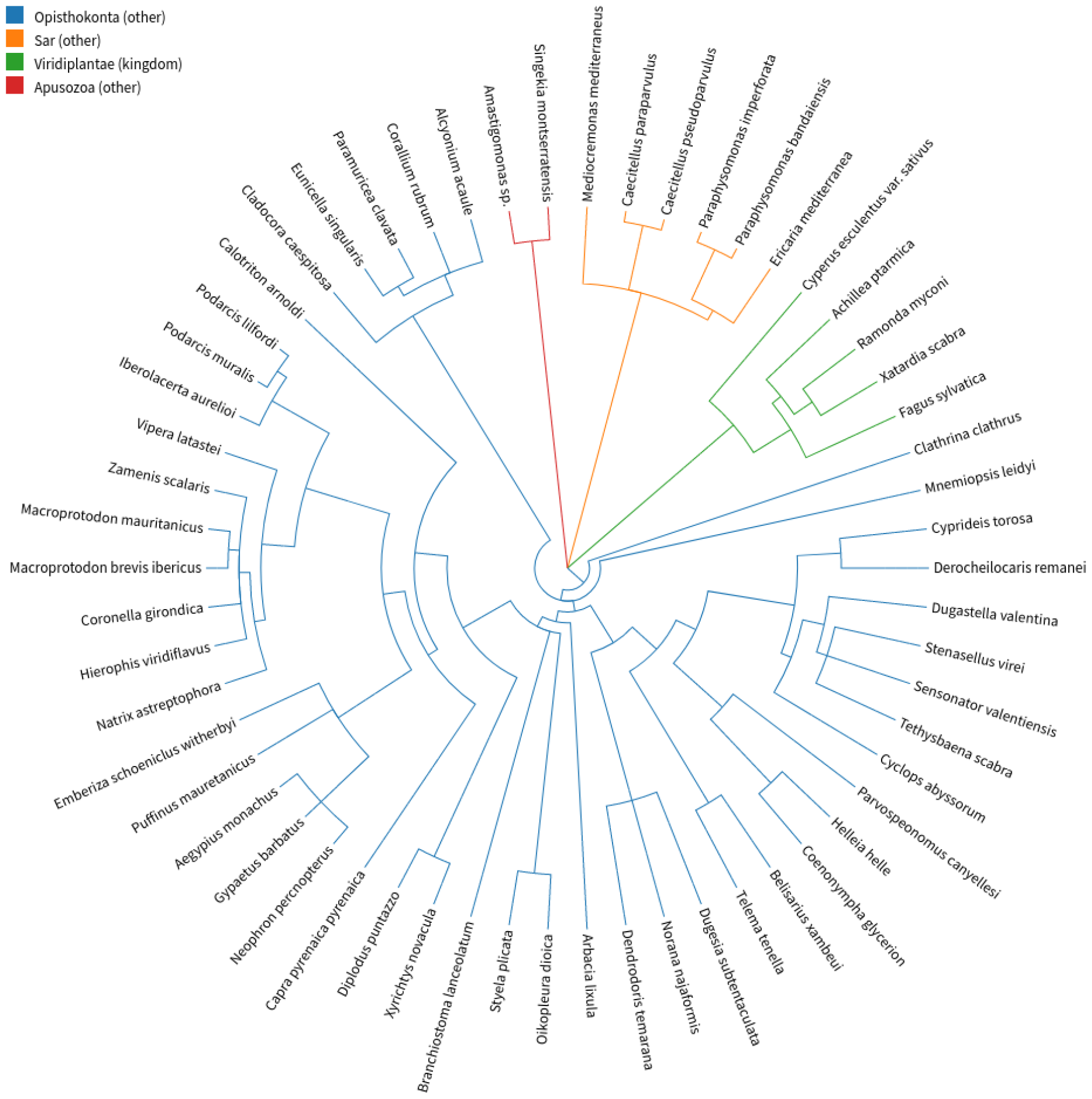
Taxonomic tree of all the 58 species currently being targeted by the CBP (as of november 2023).

One of the unique features tailored for this instance is the integration of region-specific taxonomic information, focusing on the biodiversity found in Catalonia. This customized taxonomic framework allows researchers to efficiently navigate and access relevant data on the local fauna and flora, facilitating more targeted research efforts (**Figure 5**). This use case illustrates the ability of the web portal to address region-specific research needs and its utility to foster collaboration within local scientific communities.

**Figure 5.**
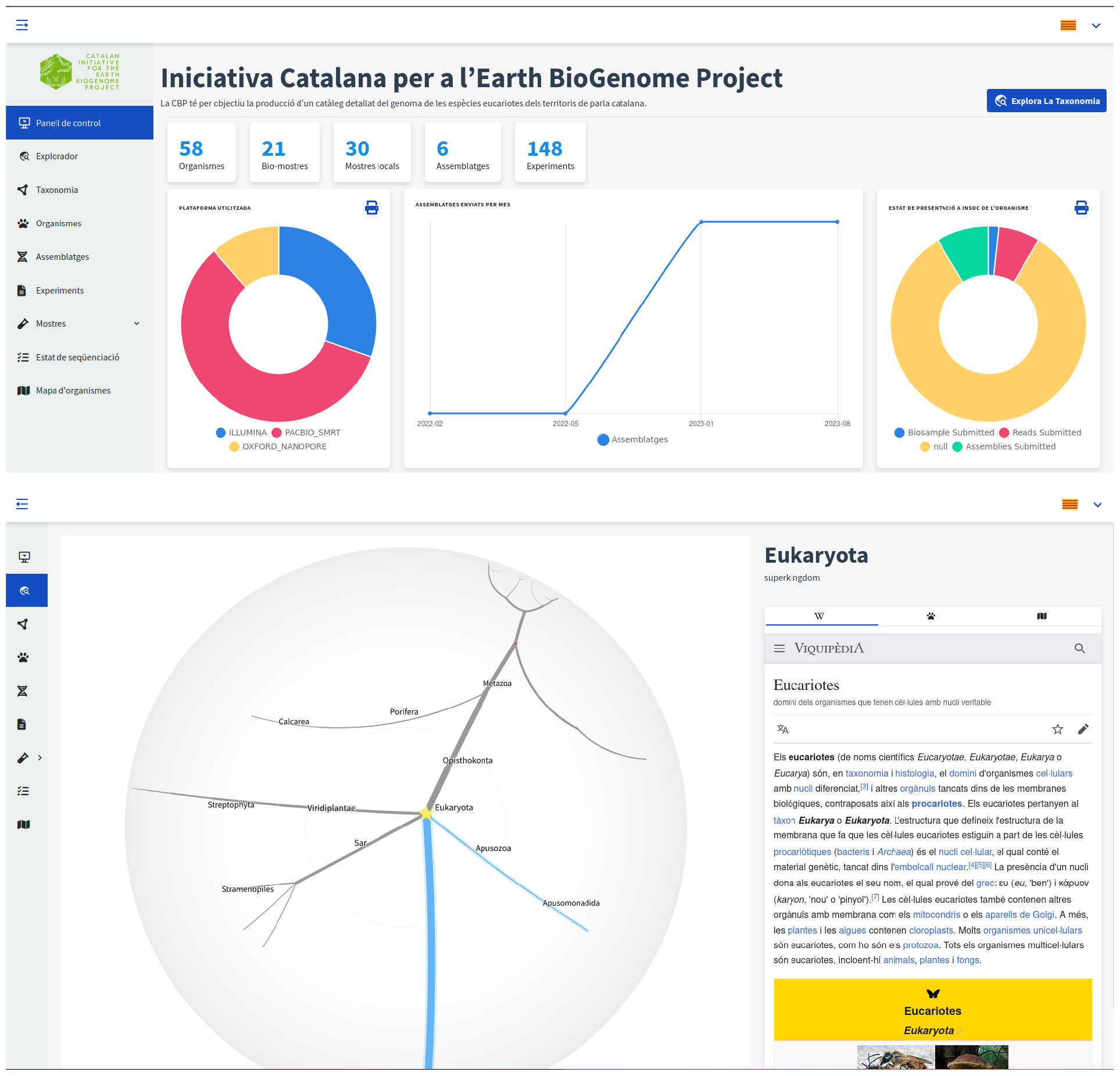
Top: Landing page of the CBP instance. Bottom: Taxonomy explorer

### EBP Instance

A dedicated BGP’ s instance tracking all the data published to INSDC under the EBP umbrella can be found at: https://ebp.biogenoma.cat/. This portal is open to users that want to add links and other relevant data not present in INSDC to the sequenced species currently targeted by the EBP. The settings of this instance are different from those in the CBP’ s instance in terms of layout, cronjobs and environment variable configuration **(Figure 6**).

**Figure 6.**
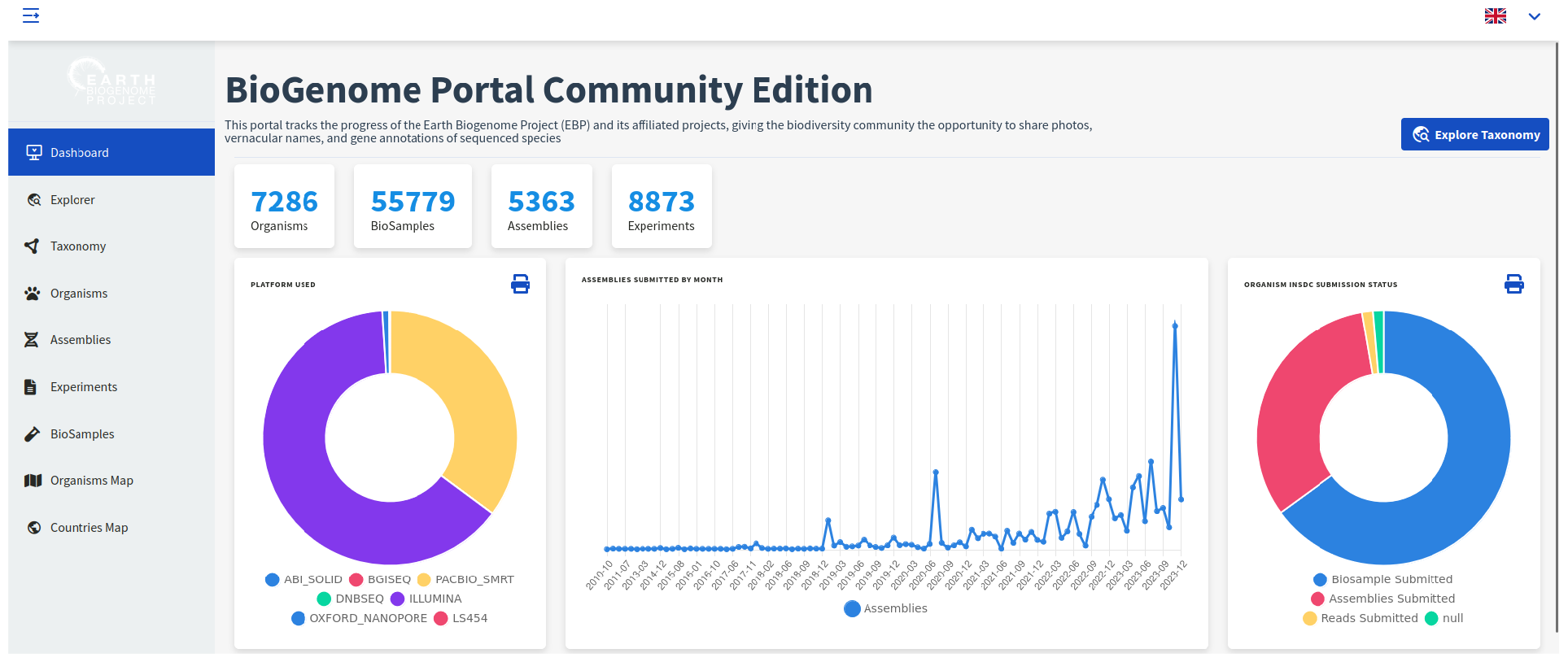
Landing page of the EBP instance

The BGP-EBP tracks all the data under the EBP bioproject accession (PRJNA533106); It contains a 3D world map showing all the species collected within the boundaries of each country (**Figure 7**) and retrieves all the biosamples submitted by under the EBP umbrella, even those not yet associated to the EBP, such as the public biosamples which project name field in their metadata matches one of the following values: ERGA, CBP, VGP or DTOL.

**Figure 7.**
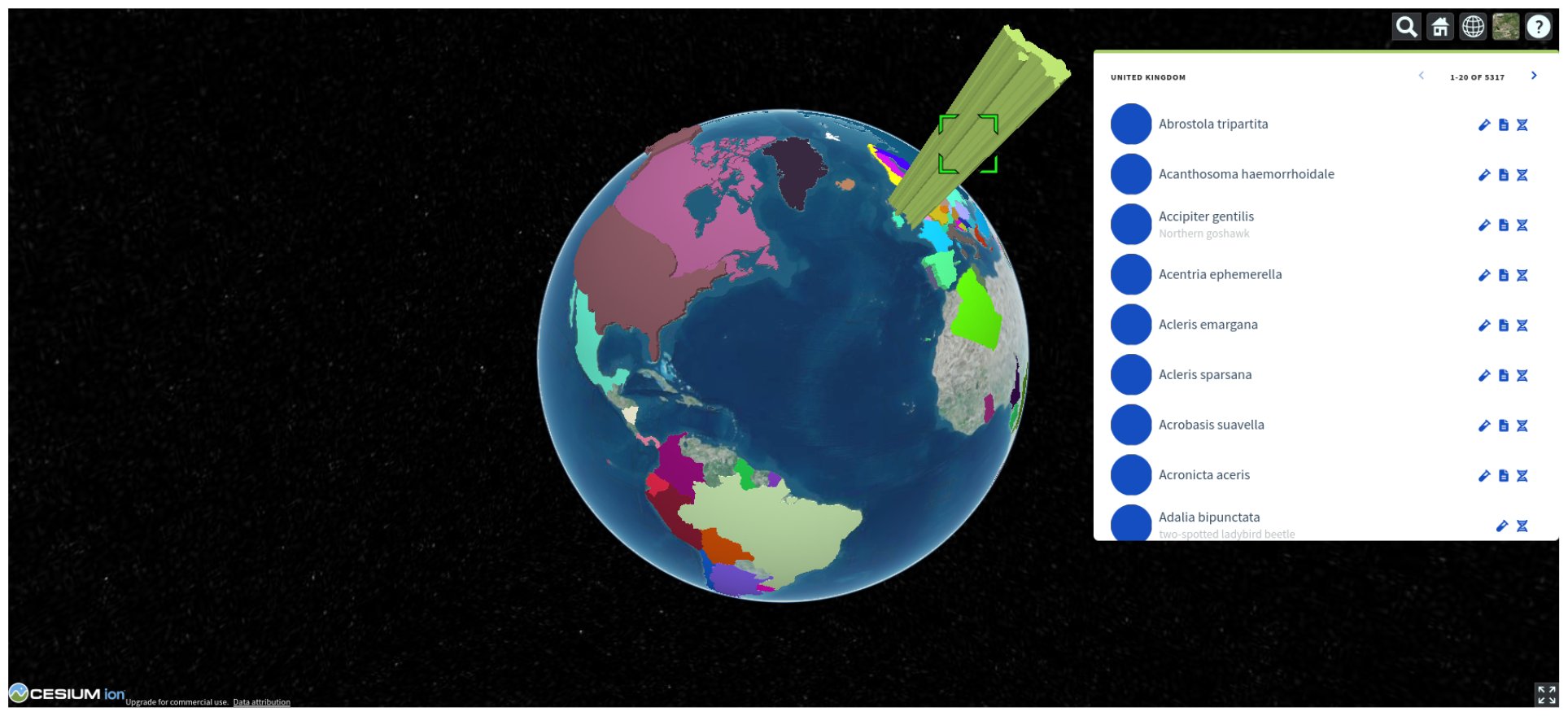
3D world map displaying the number of species collected within the boundaries of each country.

## DISCUSSION

The BioGenome Portal has been initially developed to meet the needs of the CBP, and the status of this EBP project can be accessed through an instance of the portal (https://dades.biogenoma.cat/.) The BGP-CBP portal is helping to coordinate the sequencing efforts within the CBP and facilitates data and metadata deposition in the designated international data resources. We believe that the portal will play a role in catalyzing collaborative projects between different research institutions affiliated to the CBP, and to extend the impact of the project beyond academic circles, engaging a wider community in biodiversity conservation and research initiatives.

A genuine focus of the bioinformatics developments within the CBP is in equity. In the framework of biodiversity genomics projects, equity is often understood as empowering researchers world-wide to produce data within the natural geographical region where the samples have been acquired. While the need for standardized analysis (mostly, gene annotations) across the tree of life may argue for centralized resources, bioinformatic pipelines do not need to be necessarily attached to specific sites. Developments in containerization software makes it possible to implement analysis pipelines producing highly reproducible results. These would contribute to democratization not only of data production, but also of data analysis (17).

The CBP will thus pro-active engage in promoting the development, implementation and use of light-weigth, autonomous and reproducible software. We believe that the BioGenome portal is an example of such software. It does not aim to compete with the designated EBP portal (https://www.ebi.ac.uk/biodiversity/) It is rather a generic framework for biodiversity genomics projects. It can be installed and run at any site. It can be customized to show, in addition to the data in public repositories, data specific to the site, such as vernacular names, species geographical distribution, days of sightings, etc. Sites could be other EBP nodes, but also countries, regions, shires, cities, neighborhoods, zoological parks, botanical gardens, museums, taxonomic groups, etc. Here we have specifically demonstrated the utility of the portal to display the status of both regional and taxonomic biodiversity genomics projects.

With the appropriate customization, the portal can be used as an outreach tool helping to engage the society in the understanding of biodiversity, and, in particular, of the link between genomics and biodiversity. Genomics underlines the unity of life. Understanding this, promotes the perception that biodiversity is not something separate from humans, but that humans are an inextricable part of biodiversity, and that threats to biodiversity are threats to human life on Earth.

## ACKNOWLEDGMENTS

We thank Emilio Palumbo and Ferriol Calvet, and the members of Catalan Initiative for the Earth Biogenome Project for their very useful feedback. Core support for the research at the CRG is from the CERCA Programme / Generalitat de Catalunya and from the Spanish Ministry of Science and Innovation to the EMBL partnership, Centro de Excelencia Severo Ochoa.

## SUPPLEMENTARY FIGURES

**Figure S1.**
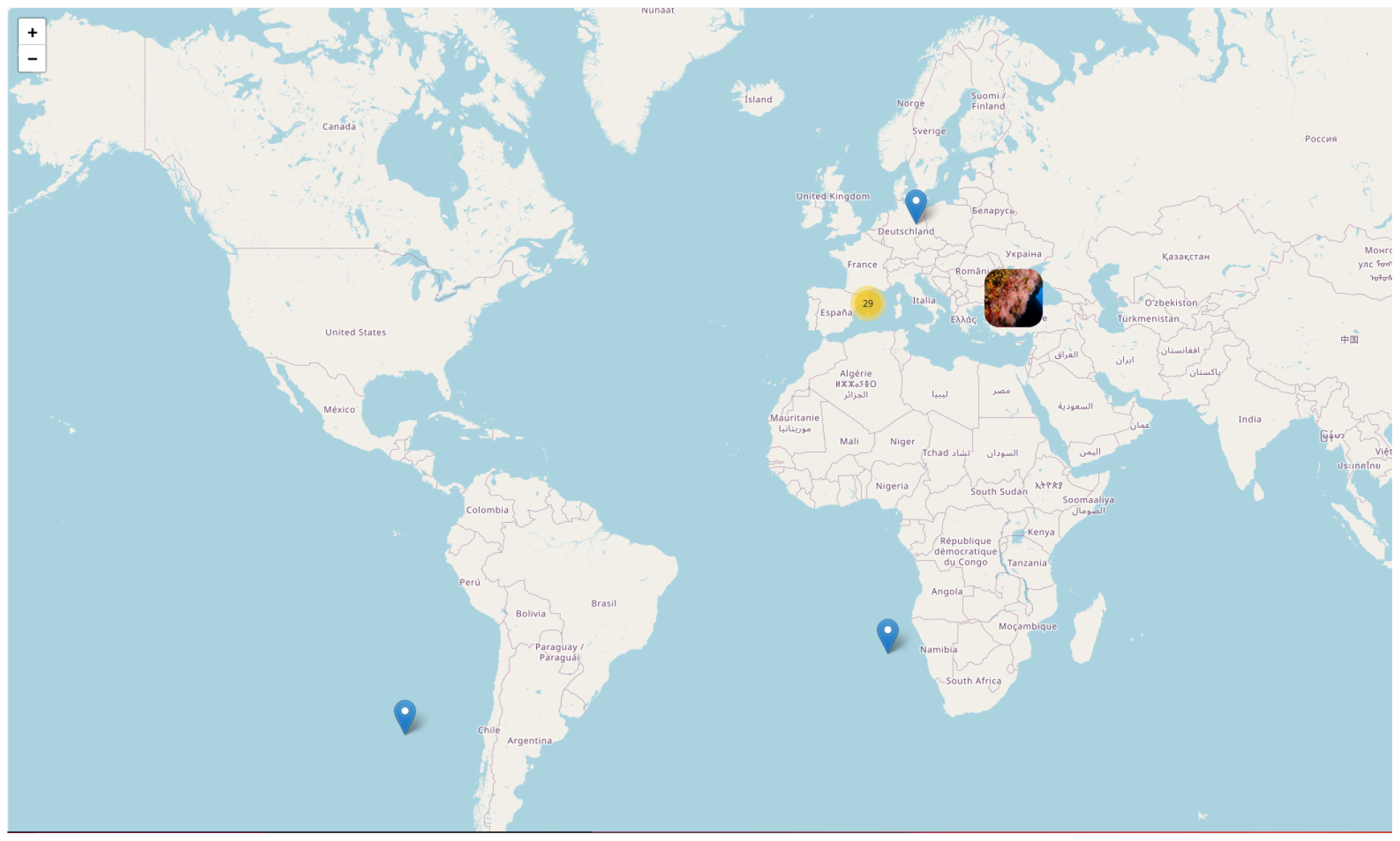
Geolocalization of the species collected by the CBP as seen through the BioGenome Portal.

**Figure S2.**
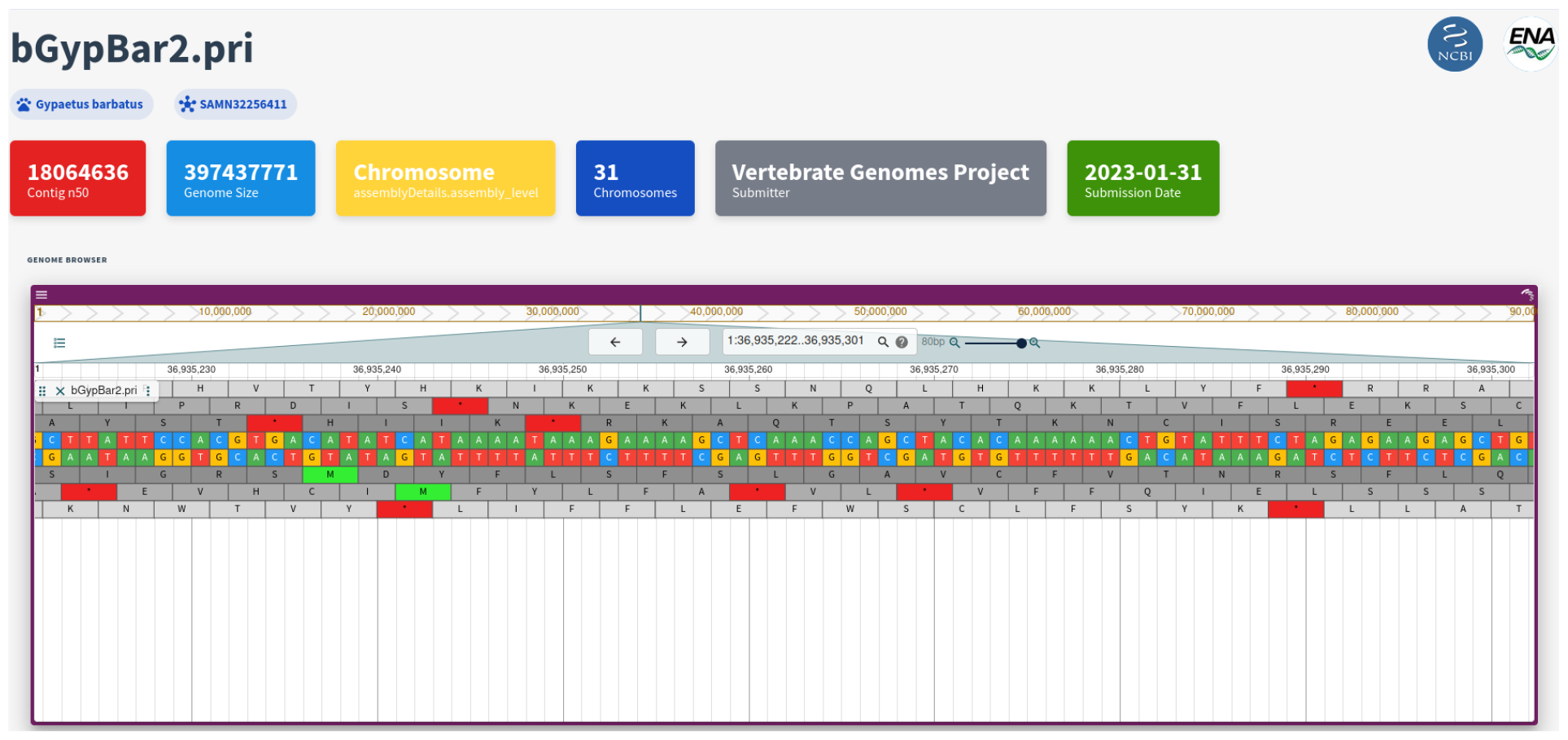
Genomic sequence of *Gypaetus barbatus*, a species sequenced by the CBP, as seen through the BioGenome Portal

